# Pre-crastination: time uncertainty increases walking effort

**DOI:** 10.1101/2020.07.17.208140

**Authors:** E Hong Tiew, Nidhi Seethapathi, Manoj Srinivasan

## Abstract

In many circumstances, humans walk in a manner that approximately minimizes energy cost. Here, we performed human subject experiments to examine how having a time constraint affects the speeds at which humans walk. First, we measured subjects’ preferred walking speeds to travel a given distance in the absence of any time constraints. Then, we asked subjects to travel the same distance under different time constraints. That is, they had to travel the given distance within the time duration provided – they can arrive early, but not late. Under these constraints, subjects systematically arrived earlier than necessary. Surprisingly, even when the time duration provided was large enough to walk at their unconstrained preferred speeds, subjects walked systematically faster than their unconstrained preferred speed. We propose that this faster-than-energy optimal speeds may be due to human uncertainty in time estimation. We show that a model assuming that humans perform stochastic optimal feedback control to arrive on time with high probability while minimizing expected energy costs predicts walking speeds higher than energy optimal, as observed in experiment.

## 1 Introduction

Humans move in a manner that at least approximately minimizes energy cost, for instance, while walking [1], running [2], sideways walking [3], walking with exoskeletons [4], walking with different step frequency, speed, and step length constraints [5], walking for short distances [6], and while choosing between walking and running under time and distance constraints [7]. When asked to walk in a preferred manner, there is evidence that humans walk close to the walking speed that minimizes the cost per unit distance [8, 9]. This tendency to walk at the energy optimal speed has been seen also in a variety of contexts, for instance, on slopes, when walking sideways, and even when walking with Down’s syndrome [3, 10]. Further, when asked to walk shorter distances, humans walk systematically slower, a behavior that also minimizes the cost of walking, including the additional cost of starting from rest and going back to rest [6].

However, there is also other evidence that suggests that humans may walk systematically faster than optimal when they perceive a cost for time, given the prospect of reward [11–13]. For instance, people in different cities and countries seem to walk at slightly different speeds [14, 15], although some of this is apparently explained by city-wise age distributions.

In this article, we measure speeds of people walking under different time constraints, and examine how these walking speeds compare with preferred walking speeds in the absence of those time constraints. We show that people walk systematically faster in the presence of time constraints, even though they would arrive on time (typically) if they ignored the time constraint and walked at their preferred speed. We then propose how this behavior could arise out of stochastic optimization, given human uncertainty in time estimations.

## 2 Methods

### Experimental methods

A total of *N* = 25 subjects participated with informed consent. We had two versions of the experiment, one performed indoors (*N*_indoor_ = 15, 9 male and 6 female, 22.2 *±* 1.26 years, height 1.68 *±* 0.08m, mass 69.71 *±* 14.06 kg, mean *±* s.d.) and one performed outdoors (*N*_outdoor_ = 10, 7 male and 3 female, 23.8 *±* 3.6 years, height 1.71 *±* 0.036 m, mass 69.72 *±* 13.17 kg, mean *±* s.d.).

Subjects walked a distance *D* = 50 m in all indoor trials and and a distance *D* = 100 m in all outdoor trials. First, each subject was asked to walk the distance *D* at a comfortable speed to compute their respective preferred walking speed. No other instructions were given. Three such preferred walking trials were performed, and the average preferred walking speed (*v*_pref_) across these three trials was computed for each subject.

Next, we performed time-constrained walking trials, where the time-constraints were based on the measured preferred speeds. In these time-constrained trials, subjects were given a stopwatch that counted down to zero from some given time *T* and were instructed to walk the distance *D*, arriving before or as the stop-watch reached zero (Figure 1a). That is, the subjects can arrive earlier than the time runs out, but not later. For the outdoor trials, subjects were instrumented with a GPS tracker (VBOX Inc.) to track their time-varying velocity during the trials.

**Figure 1:**
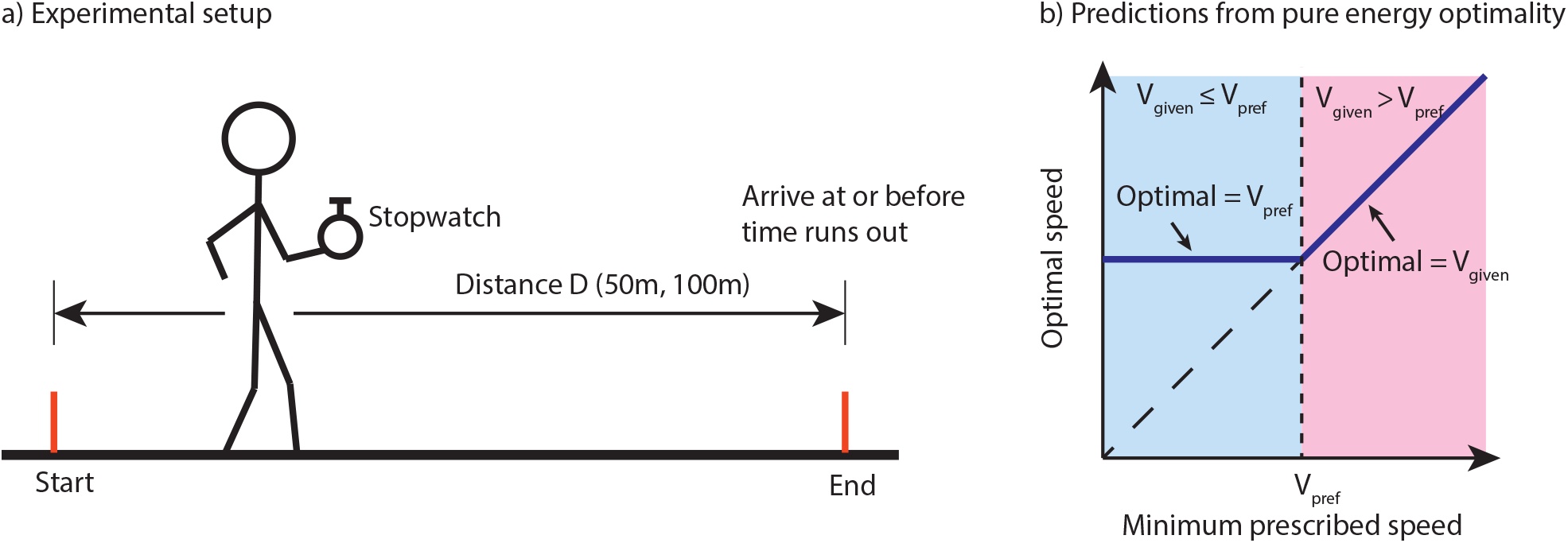
a) Subjects were asked to travel a given distance within a given amount of time. They carried a stop-watch to indicate the time remaining to complete the task. Each subject performed seven time-constrained trials, so that the minimum speed is constrained to be slower, faster, or equal to their unconstrained naturally preferred speeds. b) Predictions of pure energy optimality in locomotion, in the presence of time constraints.

For the time constrained trials, we considered seven different time limits *T*, each time limit corresponding to a minimum prescribed speed *v*_giv_; that is, *T* = *D/v*_giv_. We considered the following seven minimum speeds *v*_giv_, three slower than, three faster than, and one equal to the preferred speed: that is, *v*_pref_ 0.3 ms^*−*1^, *v*_pref_ 0.2 ms^*−*1^, *v*_pref_ 0.1 ms^*−*1^, *v*_pref_, *v*_pref_ + 0.1 ms^*−*1^, *v*_pref_ + 0.2 ms^*−*1^, *v*_pref_ + 0.3 ms^*−*1^. A total of 21 time-constrained trials were performed per subject with each time limit performed 3 times, all in random order. For all trials, we measured the total time used by the subject to traverse the distance and computed the average speed as distance over time.

### Theory: Predictions from pure energy optimality

We compare our subjects’ walking behavior in the time-constrained trials with our expectation from energy optimal walking — that is minimizing the total energy cost for the task [6, 8]. If the subjects were energy optimal, their preferred speeds in the absence of any time constraints would also be energy optimal, i.e., *v*_opt_ = *v*_pref_. In the presence of time constraints, their energy optimal speeds would depend on whether the minimum prescribed speed *v*_giv_ is greater than or less than the preferred speed *v*_pref_ (Figure 1b). When *v*_giv_ *≤ v*_pref_, that is, when there is enough time to walk at the preferred speed, the energy optimal speed is still the unconstrained preferred speed *v*_opt_ = *v*_pref_. When *v*_giv_ *> v*_pref_, that is, when walking at the unconstrained preferred speed will violate the time constraint, the energy optimal speed is the minimum speed consistent with the time constraint, namely, *v*_opt_ = *v*_giv_.

### Background: Uncertainty in time interval estimation

Executing pure energy optimality requires perfect information and error free sensory and motor processing. Humans are imperfect at estimating time without a clock or without counting or without doing something rhythmic. A number of studies have characterized human uncertainty and variability in interval timing in humans and animals, using a few different protocols e.g., see [16]. Ratikin et al [16] show that human error in estimating when a certain given time duration has elapsed increases with the time duration: in particular, the error is such that the standard deviation *σ* in the intervals estimated by the human increases linearly with the mean duration *μ*, with the ratio equal to about 0.15 (called the Weber ratio). That is, *σ* ≈ 0.15*μ*. We use this model of time uncertainty in the following. While Ratikin et al [16] display a probability distribution with compact support for the human time uncertainty (reproduced in Figure 3a), here, as a first model we use a Gaussian approximation, which is a good approximation overall, except for having infinite support. In the next version of the article, we may use a compact support approximation of the empirical distribution of time uncertainty from [16].

### Theory: Stochastic optimization and stochastic optimal control

As an alternative to pure energy optimality, we hypothesize that humans minimize the expected value of the energy cost, subject to the constraint that they arrive on time with high probability, despite any uncertainties. For the figures in this version of the manuscript, we arbitrarily set this high probability to 99%. (If we used a time-uncertainty model with compact support, we can set the probability to 100%, but this is not meaningful with a Gaussian uncertainty model.) We consider two simple models of how the human selects the speed to solve this stochastic optimization problem. In the **first model**, we assume that the human adopts an ‘open loop’ strategy: pick a speed initially and walk at that constant speed throughout. Specifically, when the human sees the time provided on the clock, this time duration is interpreted system imperfectly, that is, subject to the aforementioned time uncertainty (Figure 3a). Based on this imperfectly interpreted time duration, the human decides on a speed, which when traveled at, will minimize energy cost subject to arrival on time with 99% probability. In the **second model**, we assume that the human adopts a ‘feedback control’ strategy, in which, speed is changed multiple times through the trial. Here, the human looks at the watch, say, *N* times over the distance traveled; upon each such look at the watch, the time duration noted by the watch is interpreted by the cognitive system imperfectly, again subject to the same time uncertainty [16]. Based on this imperfectly interpreted time duration, the human decides on the optimal speed based on the same criterion as in the first model: that is, picks a speed that minimizes expected value of the energy cost while ensuring 99% probability of arrival on time. The human maintains this speed until they look at the watch again and use the new information to recompute the optimal speed. Because of the repeated optimization toward the goal based on the current state, this stochastic optimal feedback control strategy is rather like a ‘stochastic model predictive control’. The first model is a special case of the second model with *N* = 1.

In the above calculations, the expected value cost is computed as the average over possible realizations of the true time duration, given the imperfectly interpreted time duration, assuming that the nervous system has a good model of its own time uncertainty. The relevant expected value of energy and arrival probabilities are computed for each candidate walking speed by sampling from the probability distribution of time uncertainty, and then solving the stochastic optimization is simply picking the speed with the least cost, while satisfying the arrival probability constraint. We will provide a mathematical equation-based description of these calculations in a supplementary appendix in the next version of this manuscript.

## 3 Results

### Humans were able to satisfy the prescribed time constraints

When not time-constrained, our subjects had a preferred average speed of 1.53 *±* 0.22 ms^*−*1^ (mean *±* s.d.). In the time constrained trials, all subjects were able to travel the distance before the time was up, except for subject 3 in trial *n*_outdoor,7_ who arrived 3.75 s late (Figure 2a,c). That is, their average speeds were generally greater than the minimum prescribed speed *v*_giv_. In addition, not only were subjects’ averages speeds higher than *v*_giv_, but their GPS-measured time-varying speeds were also generally higher than the minimum prescribed speed *v*_giv_. This is shown in Figure 2a, displaying a histogram of percentage differences between the actual GPS-measured speed and the minimum prescribed (maximum *p* value across the 7 values = 0.333 x 10^*−*5^).

**Figure 2:**
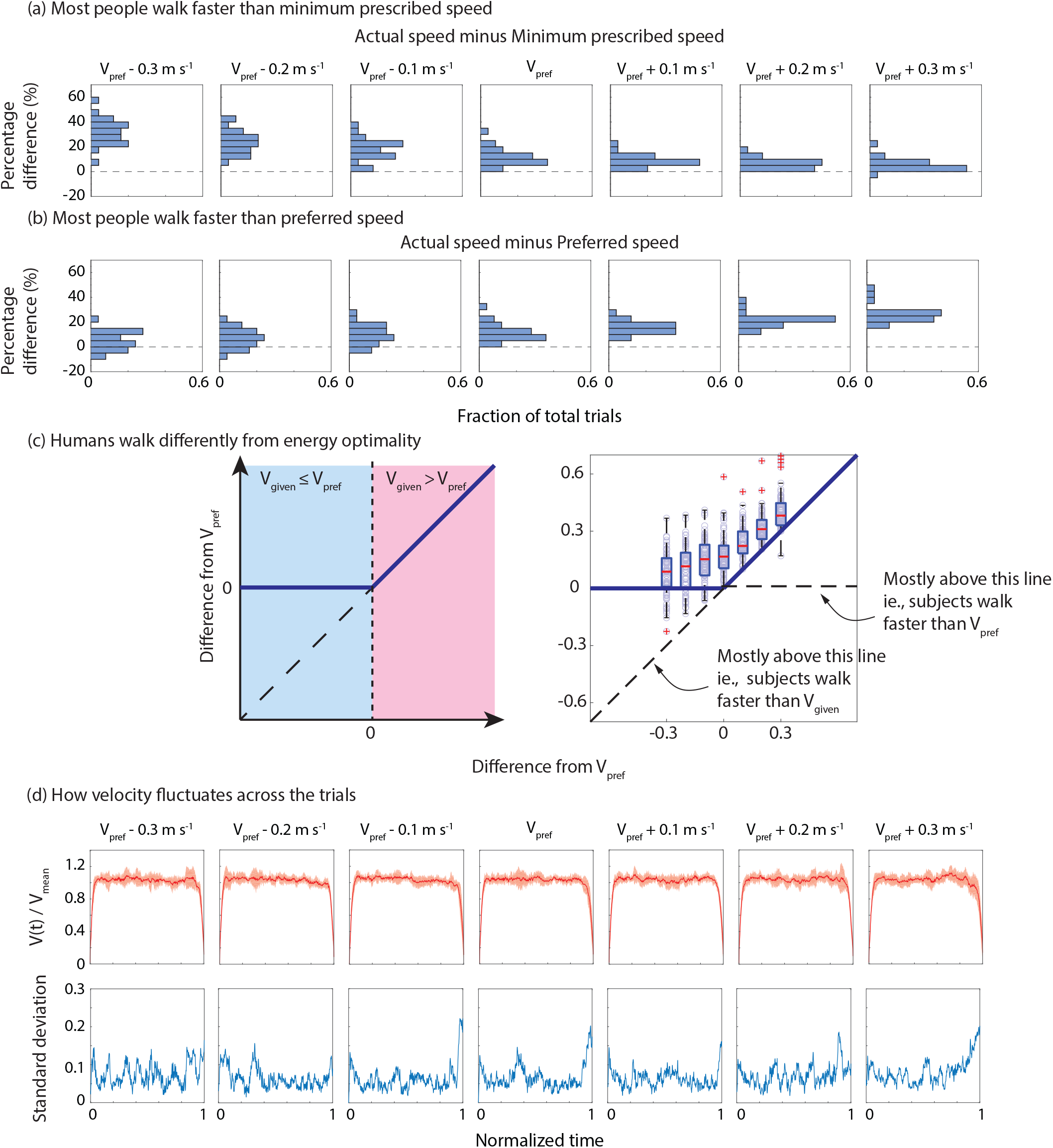
a) Percentage difference between actual average speed and minimum prescribed speed (given speed) for the seven time-constrained trials. Humans walked faster than the minimum prescribed speed, as constrained by experiment. b) Percentage difference between actual average speed and naturally preferred speed for the seven time-constrained trials. Surprisingly, humans walked faster than their naturally preferred speed, even when not constrained by experiment. c) The graph on the left shows the energy optimality prediction of how people would walk when minimizing energy cost, repeated from Figure 1b, albeit with shifted origin. The right panel shows the raw differences between walking speed in the time constrained trials and respective preferred speeds; pooled from both indoor and outdoor trials. People seem to walk faster than energy optimal. d) Normalized velocity versus normalized time, pooled across subjects in the time-constrained trials. No systematic pattern was found except for an initial acceleration and final deceleration phase. The between-subject variability in speed is higher at the end of the walking bouts, suggesting that deceleration phases were more different.

**Figure 3:**
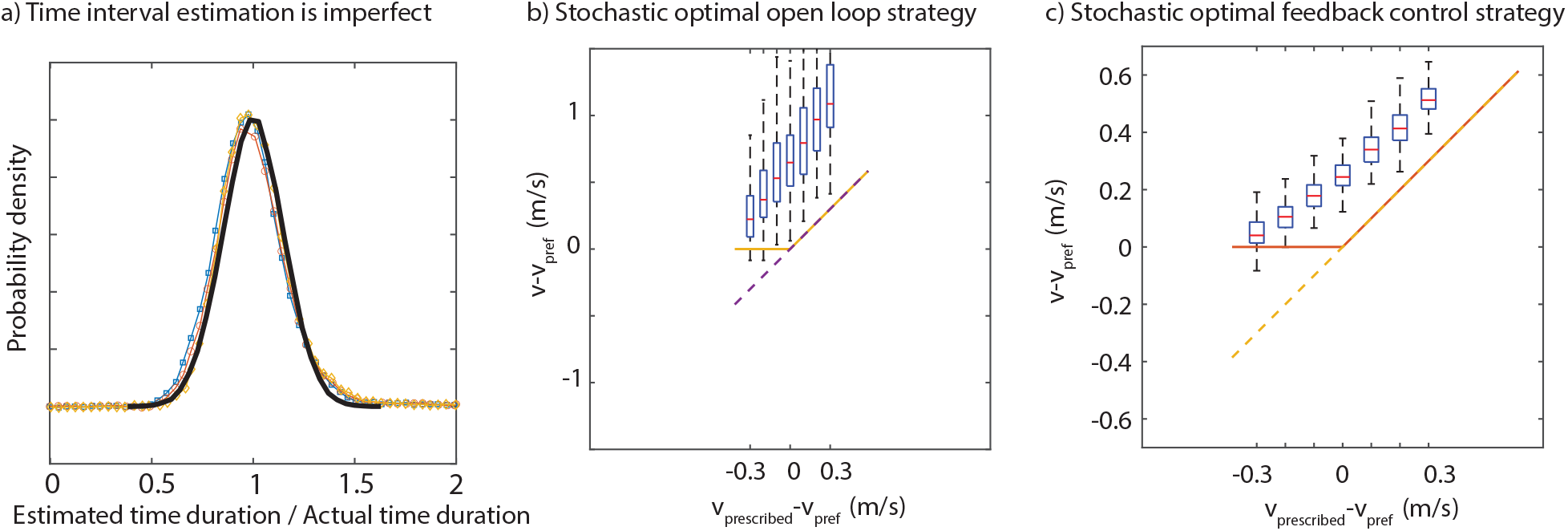
Stochastic optimization, accounting for timing uncertainty. a) Human time interval estimation is imperfect. The standard deviation of variability scales with mean time duration. Normalized by the mean, the probability distribution of estimated time becomes invariant to the mean duration. Shown are probability density data from [16] (color) and the Gaussian approximation to them (black) with Weber ratio 0.15. b) Distribution of speeds predicted by the first model, which assumes constant speed through the trial. c) Distribution of speeds predicted by the second model, which assumes stochastic optimal feedback control.

### Humans walk systematically faster than their unconstrained preferred speed

Figures 2b-c show that humans walk systematically faster than their unconstrained preferred speed *v*_pref_. For the three time-constrained cases slower than *v*_pref_, 72%, 80%, and 88% of subject walk faster than the preferred speed respectively. For the four three time-constrained cases with minimum speeds greater than equal to *v*_pref_, 100% of subjects walked faster than their preferred speeds. Overall, people walk faster than their preferred in the 160 out of the 175 time-constrained trial averages (91.43% of trials). Performing a t-test for each of seven time-limit case shows that the percent difference from *v*_pref_ is greater than zero (maximum *p* value among the seven is 0.703 x 10^*−*3^).

### Humans walk systematically differently from energy optimal

The corollary of the last two paragraphs is that humans walk systematically differently from energy optimal speeds. Further, if we assume that *v*_pref_ is optimal for the unconstrained walking trial, then the subjects walked systematically faster than energy optimal during the time-constrained trials (Figure 2c). For *v*_giv_ *<*= *v*_pref_, subjects often walked faster than *v*_pref_, even though they could walk at *v*_pref_ and comfortably satisfy the time constraint. Similarly, for *v*_giv_ *> v*_pref_, subjects walked faster than *v*_giv_, even though they could walk at *v*_giv_ to satisfy the time constraint.

### Higher variation at the beginning and end of walk

At least visually, we did not see a strong systematic pattern to the velocity fluctuations during the trials (Figure 2d), consistent across all trials. The between-subject standard deviations of velocity are higher at the beginning and at the end of the walking bouts. This suggests that between-subject acceleration/deceleration phases are different. In other words, people start and slow down at slightly different rates. When we plot the velocity fluctuation with respect to time normalized by prescribed time, we can see that there is an even much higher variation at the end of a trial (refer supplementary figure S1); this is an indication of different subjects ending their walks at different times relative to the total given time.

### No systematic differences in indoor vs outdoor

Subjects behave largely similarly in indoor and outdoor settings. Speed differences in indoor vs outdoor settings have similar distributions when compared to given and preferred speeds (supplementary figure S2 and figure S3). A two-sample t-test also showed that there is no systematic differences between both settings (minimum p-value = 0.087).

Despite no significant difference between walking indoors vs outdoors, when walking outdoors, most people tend walk faster than their preferred speed regardless of the prescribed speed. Out of 210 individual outdoor time-constrained trials conducted, only 9 trials (4.3%) fell below their respective preferred speeds and are only present in the slowest two time-constrained trials.

### Presence of time constraint systematically affects walking speed

We divided the time-constrained trials for each subject into two overlapping groups: (1) slower than or equal to preferred and (2) faster than preferred walking speeds. Each each of these groups, we fit a linear model to predict their speeds using *v*_giv_. Based on pure energy optimality, we expect the slope of this linear model to be zero for group-1, that is, that the subjects’ speed does not depend on *v*_giv_. However, we find a positive slope for all but one subject, equal to 0.318 *±* 0.365 indoors and 0.372 *±* 0.184 outdoors (mean *±* std)). This suggests that subjects increase their speeds systematically as *v*_giv_ increases, rather than just walk at *v*_pref_.

For group-2, from energy optimality, we expect the slope with respect to be equal to one, that is people walk at *v*_giv_. Instead, slope is 0.761 *±* 0.219 for indoors and 0.729 *±* 0.166 for outdoors. This slope less than one means that people walk with speeds closer to *v*_giv_ for higher speed cases, while always walking faster than *v*_giv_.

### Stochastic optimal control predicts faster speeds than purely energy optimal

We find that both stochastic optimization models result in a distribution of mean walking speeds that are systematically faster than the energy optimal (Fig 3b-c). The first model predicts average speeds that are much higher than energy optimal, primarily because if the speed needs to be constant, one needs a high speed to mainly a high probability of on time arrival (Fig 3b). The second model, which allows changing the speed multiple times over the walking bout based on current time duration remaining, also produces higher than energy optimal speeds on average, but not as much higher as the first model (Fig 3c). Defending against time uncertainty implies that humans should arrive earlier than the prescribed speed – but that they also arrive earlier than purely energy optimal speeds for low prescribed speeds is surprising and suggests that the effect of time uncertainty is quite large. We notice that the predicted speed grows with slope less than one with respect to the prescribed speed, as also see in data.

## 4 Discussion

The results of our experiment could be viewed as a special case of a recently named phenomenon, precrastination. Pre-crastination is the tendency to complete or begin a task as soon as possible at the expense of extra physical effort [17], possibly to reduce cognitive load. In our experiments, it is possible for subjects make the task of arriving before the deadline ‘simpler’ by walking faster than energy optimal.

While we have argued that subjects walk differently from energy optimal, it is still the case that our subjects are still close to their energy optimal speeds compared to all the speeds that they are capable of (for instance, running). Thus, our result could be considered a small modification of energy optimality under certain situations, rather than ‘refuting’ energy optimality.

Indeed, we have shown that model the subjects’ behavior as being optimal in a stochastic or robust control sense may explain why people arrive systematically earlier. A more detailed mathematical treatment of the modeling and the optimization will be presented in a later version. Here, we have explained why humans systematically arriving earlier by accounting for the subjects’ uncertainties in their estimation of time alone, but we might also consider uncertainties in distance, imperfections in their control of velocity, and their motivation to not arrive late (as the task demanded). The detailed feedback control mechanism that governs how subjects arrive on time could also identified from the data in future versions of this work.

Energy optimality has usually been assumed, perhaps only implicitly, as a simple first theory and not something that is likely to provide a perfect prediction of human behavior in every situation. Given the broad success of energy optimality as a predictive theory, understanding even small systematic deviations from it are useful. This article has provided one such situation, in which we argue that humans may expend more energy and walk at higher speeds to defend against time uncertainly. Of course, our model is still based on energy optimality, except accounting for time uncertainty and reframing optimality in a stochastic expected value sense in contrast to deterministic energy optimality, which may superficially appear to be not energy optimal.

## Ethics statement

All subjects participated with informed consent. The experiments were approved by the Ohio State University Institutional Review Board.

## Notes

### Competing Interest Statement

The authors have declared no competing interest.

### Summary of Updates

We have added a new mathematical model relying on stochastic optimal control that explains the data in the previous manuscript version.

## References

[1] Maxwell Donelan J, Kram R, Arthur D K. Mechanical and metabolic determinants of the preferred step width in human walking. Proceedings of the Royal Society of London Series B: Biological Sciences. 2001;268(1480):1985–1992.

[2] Högberg P. How do stride length and stride frequency influence the energy-output during running? European journal of applied physiology and occupational physiology. 1952;14(6):437–441.

[3] Handford ML, Srinivasan M. Sideways walking: preferred is slow, slow is optimal, and optimal is expensive. Biology letters. 2014;10(1):20131006.

[4] Selinger JC, O’Connor SM, Wong JD, Donelan JM. Humans can continuously optimize energetic cost during walking. Current Biology. 2015;25(18):2452–2456.

[5] Bertram JE, Ruina A. Multiple walking speed–frequency relations are predicted by constrained optimization. Journal of theoretical Biology. 2001;209(4):445–453.

[6] Seethapathi N, Srinivasan M. The metabolic cost of changing walking speeds is significant, implies lower optimal speeds for shorter distances, and increases daily energy estimates. Biology letters. 2015;11(9):20150486.

[7] Long III LL, Srinivasan M. Walking, running, and resting under time, distance, and average speed constraints: optimality of walk–run–rest mixtures. Journal of The Royal Society Interface. 2013;10(81):20120980.

[8] Srinivasan M. Optimal speeds for walking and running, and walking on a moving walkway. Chaos: An Interdisciplinary Journal of Nonlinear Science. 2009;19(2):026112.

[9] Ralston HJ. Energy-speed relation and optimal speed during level walking. Internationale Zeitschrift für Angewandte Physiologie Einschliesslich Arbeitsphysiologie. 1958;17(4):277–283.

[10] Agiovlasitis S, Motl RW, Ranadive SM, Fahs CA, Yan H, Echols GH, et al. Energetic optimization during over-ground walking in people with and without Down syndrome. Gait & posture. 2011;33(4):630–634.

[11] Shadmehr R, Huang HJ, Ahmed AA. A representation of effort in decision-making and motor control. Current biology. 2016;26(14):1929–1934.

[12] Yoon T, Geary RB, Ahmed AA, Shadmehr R. Control of movement vigor and decision making during foraging. Proceedings of the National Academy of Sciences. 2018;115(44):E10476–E10485.

[13] Summerside EM, Shadmehr R, Ahmed AA. Vigor of reaching movements: reward discounts the cost of effort. Journal of neurophysiology. 2018;119(6):2347–2357.

[14] Bornstein MH, Bornstein HG. The pace of life. Nature. 1976;259(5544):557.

[15] Levine RV, Norenzayan A. The pace of life in 31 countries. Journal of cross-cultural psychology. 1999;30(2):178–205.

[16] Rakitin BC, Gibbon J, Penney TB, Malapani C, Hinton SC, Meck WH. Scalar expectancy theory and peak-interval timing in humans. Journal of Experimental Psychology: Animal Behavior Processes. 1998;24(1):15.

[17] Rosenbaum DA, Gong L, Potts CA. Pre-crastination: Hastening subgoal completion at the expense of extra physical effort. Psychological Science. 2014;25(7):1487–1496.

